# ALKBH5 regulates somatic cell reprogramming in a phase specific manner

**DOI:** 10.1101/2021.05.10.443389

**Authors:** Sherif Khodeer, Arne Klungland, John Arne Dahl

**Affiliations:** Department of Microbiology, Oslo University Hospital, Rikshospitalet, Forskningsveien 1, 0373.Oslo, Norway; Department of Biosciences, Faculty of Mathematics and Natural Sciences, University of Oslo, 0316.Oslo, Norway

**Keywords:** *Alkbh5*, reprogramming, induced pluripotent stem cells (iPSCs), *Nanog*

## Abstract

Establishment of the pluripotency regulatory network in somatic cells by introducing four transcriptional factors, (Octamer binding transcription factor 4 (OCT4), SRY (sex determining region Y)-box 2 (SOX2), Kruppel - like factor 4 (KLF4), and cellular-Myelocytomatosis (c-MYC) provides a promising tool for cell-based therapies in regenerative medicine. Still, the mechanisms at play when generating induced pluripotent stem cells from somatic cells is only partly understood. Here we show that the RNA specific N6-methyladenosine (m^6^A) demethylase ALKBH5 regulates somatic cell reprogramming in a stage specific manner. Knockdown or knockout of *Alkbh5* in the early reprogramming phase impairs the reprogramming efficiency by reducing the proliferation rate through arresting the cells at G2/M phase and decreasing the mesenchymal-to-epithelial transition (MET) rate. However, there is no significant change in reprogramming efficiency when *Alkbh5* is depleted at the late phase of reprogramming. On the other hand, ALKBH5 overexpression at the earlyreprogramming phase has no significant impact on reprogramming efficiency, while overexpression at the late phase enhances the reprogramming by stabilizing *Nanog* transcripts resulting in upregulated *Nanog* expression. Our study provides mechanistic insight into the crucial dynamic role of ALKBH5 in regulating somatic cell reprogramming at the posttranscriptional level.

## Introduction

The four transcriptions factors OCT4, SOX2, KLF4, and c-MYC (OSKM) are sufficient to reprogram and induce pluripotency when ectopically expressed in mouse or human somatic cells, to generate induced pluripotent stem cells (iPSCs) [1, 2]. These reprogrammed iPSCs are highly similar to their pluripotent embryonic stem cell (ESC) counterparts in transcriptional profile and epigenetic landscape [3-5], and show infinite self-renewal capability [2], and the ability to differentiate to the three germ layers *in vivo* and *in vitro* [6]. Therefore, iPSC technology provide an ideal tool for drug screening, patient-specific disease modeling, and hold great promise for therapeutic applications in the future [7].

The early phase of the reprogramming process is characterized by stochastic events [8], in which the mesenchymal genes are downregulated, while epithelial genes are upregulated in a process known as mesenchymal-epithelial transition (MET), together with clear morphological transformation accompanied with increased proliferation rate to form cellclusters [9, 10]. Despite that, most fibroblasts exposed to iPSC reprogramming conditions fail to achieve the proper morphological changes and remain in a fibroblast like morphology. These trapped cells undergo senescence, apoptosis, and cell cycle arrest, which in turn explain the low efficiency of the reprogramming process [11-13]. Besides, several studies have demonstrated that cell cycle regulators including p21, p53 or p16/INK4A are barriers to the reprogramming process and their depletion enhances the reprogramming process [14-17].

The late phase of the reprogramming process is considered deterministic, in which reactivation of endogenous *Sox2* expression is considered a rate-limiting step for acquiring the ESCs identity [8]. This phase is also characterized by removal of somatic epigenetic memory, telomere elongation, expression of endogenous pluripotency genes, and establishment of pluripotency specific epigenetic and transcriptional profiles [9, 10].

The N6-methyladenosine (m^6^A) modification, methylation of the N6 position of the adenosine base, is the most abundant internal posttranscriptional modification in mammalian mRNA [18]. It was recently showed that m^6^A modification is reversible and its presence is regulated through coordination of several modulators [19, 20]. Positioning of m^6^A is mediated by methyl transferase-like 3 (METLL3), methyl transferase-like 14 (METLL14) and Wilms’ tumor 1-associating protein (WTAP) [21-24]. Removal of m^6^A is carried out by the demethylases fat mass and obesity-associated protein (FTO) and alkylated DNA repair protein AlkB homolog 5 (ALKBH5) [20, 25]. Furthermore, the m^6^A modification is recognized and bound by readers including YTH domain-containing proteins 1-3 (YTHDF1-3) and (YTHDC1 and 2) which in turn facilitate downstream processing such as mRNA splicing, stabilization, translation or degradation [26-28].

ALKBH5 is one of nine mammalian members of the AlkB family of Fe (II) and α-ketoglutarate dependent dioxygenases, and can demethylate the m^6^A modification in RNA to adenosine (A) [24]. We have previously shown that *Alkbh5* is highly expressed in meiotic cells of the testis and is mainly localized to the nucleus [24]. ALKBH5 has been shown to regulate various biological and pathophysiological processes including; autophagy, glioblastoma, breast cancer, lung cancer and infertility [24, 28-32]. In addition, the heterogeneity in *Alkbh5* expression in several cancer models has led to suggestions of a putative oncogenic or tumor suppressive role [33]. Despite extensive studies on ALKBH5 in different biological systems, the functional and regulatory role of ALKBH5 in somatic cell reprogramming has not been addressed. In this study, we dissected the precise role of ALKBH5 in the reprogramming process and our data revealed that ALKBH5 exhibits a biphasic role during somatic cell reprogramming. Depletion of *Alkbh5* in the very early phase of reprogramming impairs the reprogramming process through downregulation of Cyclin B1 and B2 resulting in reduction in the cell proliferation rate, and arresting cells at G2/M phase accompanied by decreasing the rate of MET. In the late phase, overexpression of Alkbh5 stabilizes *Nanog* transcripts resulting in upregulated *Nanog* expression, which in turn enhances the reprogramming efficiency.

## Results

### ALKBH5 depletion in early phase impairs reprogramming efficiency

To explore the role of ALKBH5 in reprogramming, we first examined the expression of *Alkbh5* during the reprogramming process, and we found that the expression of ALKBH5 is gradually upregulated during reprogramming at both the mRNA and protein levels (Fig. 1A, B). Then we used two different short hairpin RNAs (shRNA) to knockdown *Alkbh5* expression (Fig. 1C). As expected by knocking down *Alkbh5*, we found that the total m^6^A level at mRNA was highly increased compared to the controls (supplementary Fig.1A)

**Figure 1.**
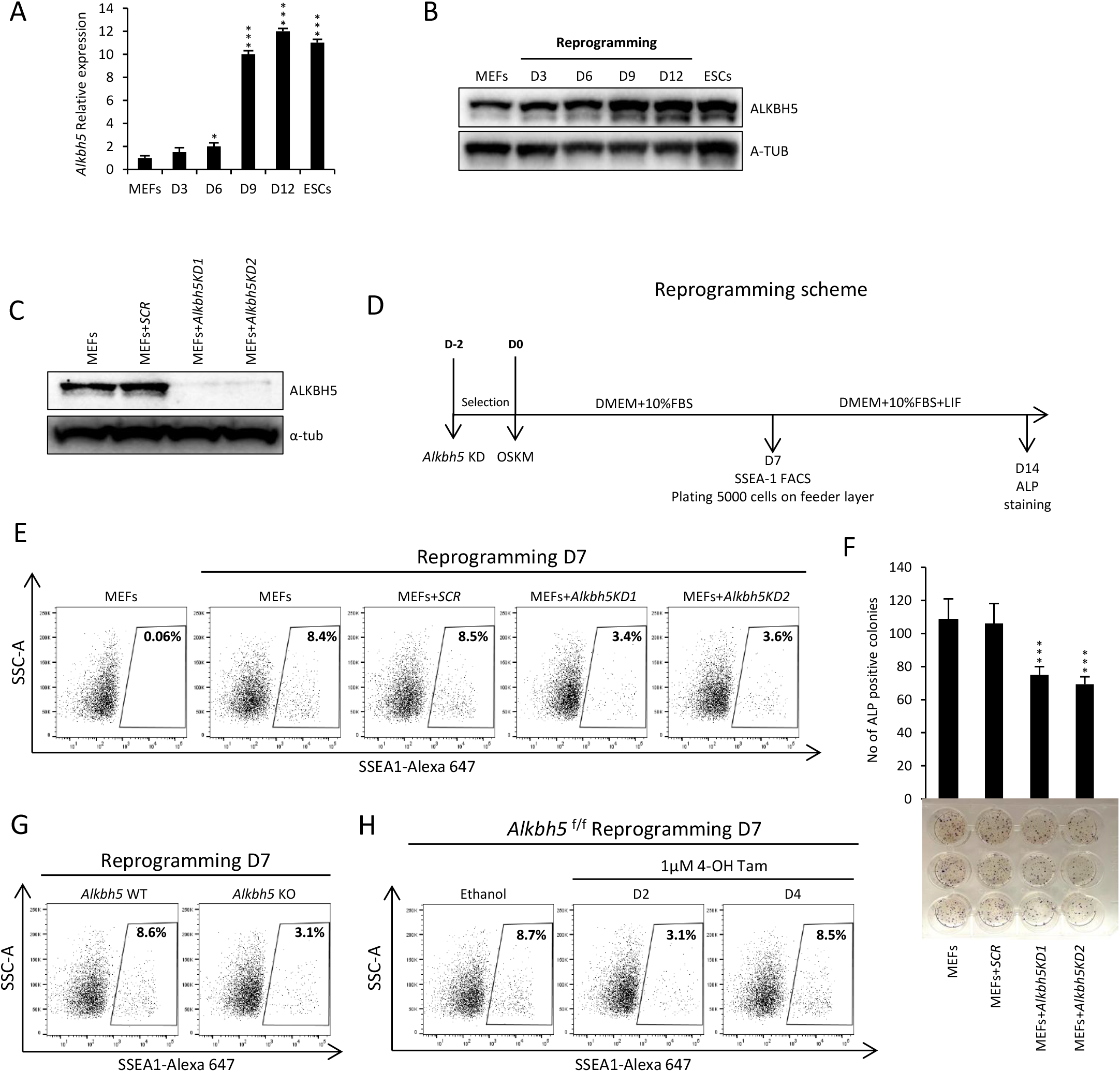
*Alkbh5* depletion impairs the somatic cell reprogramming efficiency. **(A)** Relative expression of *Alkbh5* during somatic cell reprogramming detected by qPCR. Mouse embryonic fibroblasts (MEFs) and mouse embryonic stem cells (ESCs) cultured in serum plus leukemia inhibitory factor LIF (S/L), were used as a negative and positive controls of the pluripotency, respectively. Data are normalized to the housekeeping gene glyceraldehyde-3-phosphate dehydrogenase (*Gapdh*). **(B)** Immunoblot analysis of ALKBH5 protein level during reprogramming. Alpha-tubulin (A-TUB) was used as loading control. **(C)** Immunoblot analysis of ALKBH5 protein level in MEFs after lentiviral infection with either scrambled or two different shRNAs targeting Alkbh5. (A-TUB) was used as loading control. **(D)** Experimental design showing the timing of *Alkbh5* knockdown, onset of reprogramming, SSEA1 and ALP detection. **(E)** Fraction of SSEA1 positive cells determined by FACS analysis after *Alkbh5* knockdown during the early phase of reprogramming. MEFs are used as negative control. **(F)** Reprogramming efficiency was measured by counting the number of ALP positive colonies. **(G)** Fraction of SSEA1 positive cells determined by FACS for reprogrammed wild type (WT) and knockout (KO) Alkbh5 MEFs assessed at day 7 of reprogramming. **(H)** Fraction of SSEA1 positive cells determined by FACS analysis of reprogrammed homozygous Floxed *Alkbh5* (*Alkbh5*^*f/f*^) treated with Ethanol as a control or 1µM of 4-hydroxy Tamoxifen (4-OH Tam) for depletion of *Alkbh5* at either day 2or day 4. Data are shown as mean ± SD; n = 3, *P < 0.05, **P < 0.01, and ***P < 0.001 mean ± SD deviation of triplicate samples.

Next, we sat up a reprogramming system where *Alkbh5* is knocked down 2 days before induction of retroviral reprogramming factors (OSKM) in mouse embryonic fibroblasts (MEFs), and we assessed the reprogramming efficiency on day 7 by flow cytometry and on day 14 by alkaline phosphatase (ALP) staining (Fig. 1D). Interestingly, *Alkbh5* knockdown significantly reduced the reprogramming efficiency by decreasing the percentage of stage-specific embryonic antigen1 (SSEA1) positive cells (an early reprogramming marker) on day 7, and the number of ALP positive colonies on day 14, as compared to controls (Fig. 1E, F). To substantiate these data, we derived of *Alkbh5* knockout (KO) MEFs and we found the reprogramming efficiency of *Alkbh5* (KO) MEFs is greatly reduced compared to wild type (WT) MEFs either on day 7 or day 14 by decreasing the percentage of SSEA1 positive cells in the population (Fig. 1G and supplementary Fig.1B, C) [24]. Taken together, these data suggests that *Alkbh5* depletion at the early phase of reprogramming impairs somatic cell reprogramming.

To further characterize the time specific role of ALKBH5, we took advantage of adoxycycline (DOX) inducible short hairpin RNA (shRNA) expression to suppress the expression of *Alkbh5* at specific time points during reprogramming (Supplementary Fig. 1 D). We found that *Alkbh5* knockdown at the very early stage of reprogramming, earlier than day 3, has the largest impact on reducing the reprogramming efficiency as shown by decreased fraction of SSEA1 positive cells on day 7 and 14 of reprogramming (Supplementary Fig. 1, E). On the other hand, we did not see any significant change in reprogramming efficiency when *Alkbh5* was knocked down specifically at a later time than day 3 of the reprogramming process (Supplementary Fig. 1 E, F). Furthermore, we derived homozygous floxed *Alkbh5* (*Alkbh5* ^*f/f*^) MEFs and we used a 4-hydroxy tamoxifen (4-OH Tam) inducible Cre recombinase system, in which Cre is flanked by mutated ligand-binding domains of the murine estrogen receptor (Mer-Cre-Mer), to deplete *Alkbh5* at specific time points during reprogramming (Supplementary Fig. 1 G-I) [24]. Consistent with our time specific knockdown data, depletion of *Alkbh5* only at the very early stage (day 2) of reprogramming impairs the reprogramming as measured by a decreased percentage of SSEA1 positive cells in the population (Fig. 1 H). Time specific depletion of *Alkbh5* at day 8 or 10 of reprogramming has no significant impact on the reprogramming efficiency (Supplementary Fig. 1 J). We further confirmed our data by treating homozygous floxed *Alkbh5* MEFs with (4-OH Tam) to deplete *Alkbh5* at different time points of reprogramming, and we found that only *Alkbh5* depletion on day 2 or day 4 has a major impact on reducing the reprogramming efficiency as measured by alkaline phosphates staining at day 14 (Supplementary Fig. 1 K). In conclusion, only *Alkbh5* depletion at the very early stage of reprogramming negatively affects the reprogramming process.

### Effect of *Alkbh5* removal during the early phase of reprogramming on cell cycle regulators and MET

To investigate the mechanism involved in reduced reprogramming efficiency resulting from loss of *Alkbh5*, we focused on two important events; cell proliferation and MET that have both been reported to be critical to the early phase of reprogramming [9, 10]. First, we explored the impact of *Alkbh5* removal on proliferation and apoptosis during the early phase of reprogramming. Our data revealed that *Alkbh5* knockdown during the early phase of reprogramming increases the percentage of cells at G2/M phase (Fig. 2A, B). Additionally, *Alkbh5* depletion resulted in reduced cell proliferation (Fig. 2C). However, we did not see any significant changes in the percentage of Annexin positive cells as compared to the control, indicating that the reduction in cell number is mainly due to G2/M cell cycle arrest (Supplementary Fig. 2A-C). Next, we assessed the expression of factors of the mitotic checkpoint complex (MCC) and found that Cyclin B1 and B2 are markedly downregulated at both the RNA and protein level after knocking down *Alkbh5* during the early phase of reprogramming (Fig. 2D, E). Other MCC factors such as *Cdc20, Mad1, Mad2, Bub1* and *Bub3* or G1 phase cell cycle regulators such as *p16* and *p19* were not significantly affected (Fig. 2D, E and supplementary Fig. 2D). To validate our *Alkbh5* knockdown data, we used *Alkbh5*^*f/f*^ MEFs and induced *Alkbh5* removal by (4-OH Tam) just 8 hours after reprogramming induction. In agreement with our knockdown data, we found reduction in Cyclin B1 and B2 levels showing that this phenotype is present with the loss of *Alkbh5* both in MEFs and in the early reprogramming process (Supplementary Fig. 2E). It is also noteworthy that depletion of *Alkbh5* in MEFs decreased the proliferation rate, and induced cell cycle arrest at G2/M phase accompanied by reduction in the protein level of both Cyclin B1 and B2 (Supplementary Fig. 2F - I). This is consistent with what we observed during reprogramming (Fig. 2A -E). Thereafter, we assessed the MET process at day 6 of reprogramming in which *Alkbh5*was knocked down 2 days before reprogramming induction. Our qPCR and western blot data revealed that *Alkbh5* depletion impair the MET process by decreasing the rate of downregulation of mesenchymal markers such as Platelet Derived Growth Factor Receptor Beta (*PDGFRβ*), Snail Family Zinc Finger 2 (*Slug*), Zinc Finger E-Box Binding Homeobox 1 (*Zeb1*) and Zinc Finger E-Box Binding Homeobox 2 (*Zeb2*), and upregulation epithelial markers such as E-cadherin (*E-cad*), Epithelial Cell Adhesion Molecule (*Epcam*), and *Occludin* (Fig. 2F, G). Tamoxifen induced deletion of *Alkbh5* eight hours after reprogramming supported this role of ALKBH5 in the MET process during reprogramming (Supplementary Fig. 2J). The role of ALKBH5 in MET is further supported by our observations on by tracking morphological changes during reprogramming after *Alkbh5* depletion (Supplementary Fig. 2K). In summary, *Alkbh5* is required for proper cell proliferation and for proper MET in the early phase of reprogramming.

**Figure 2.**
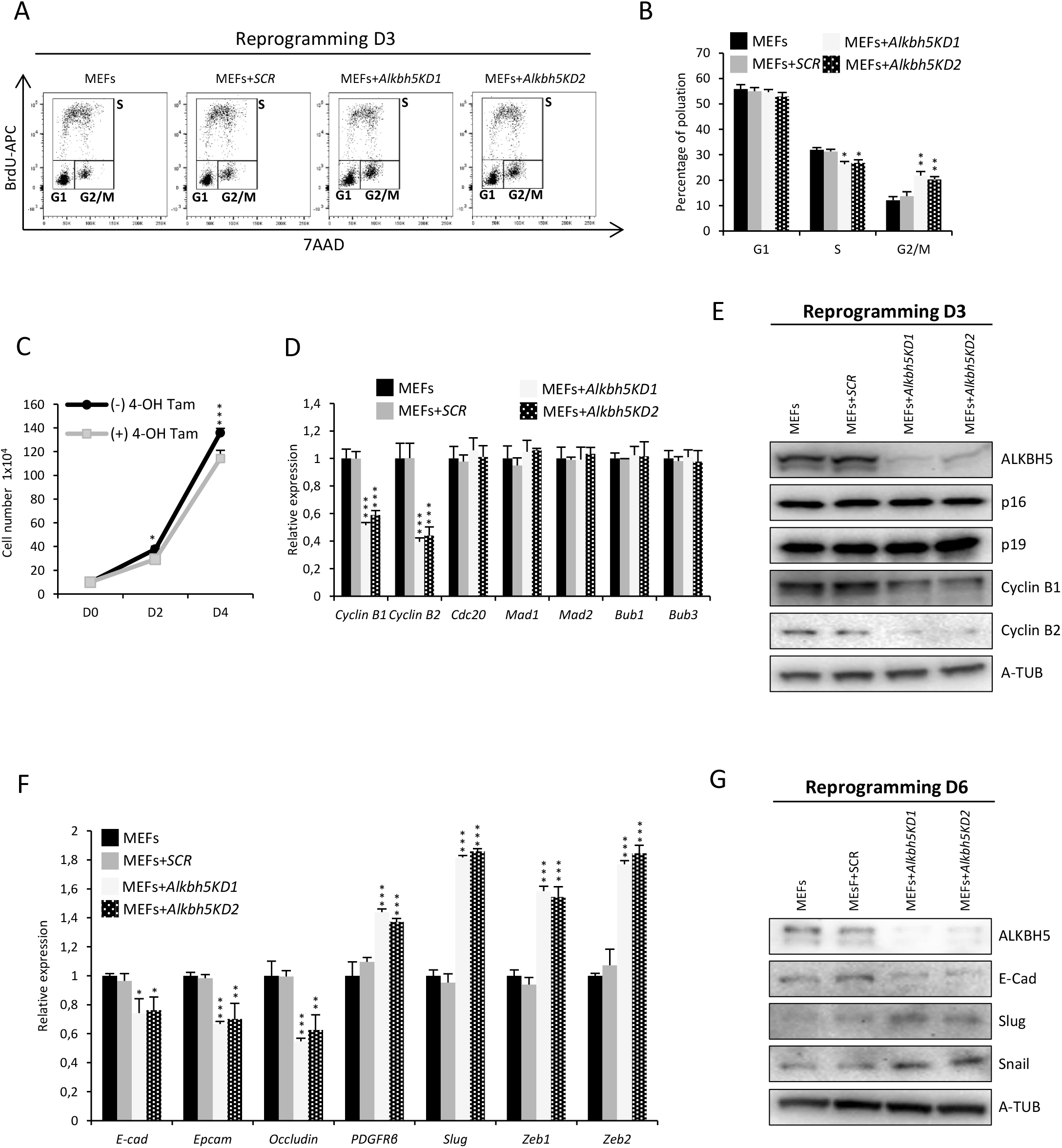
*Alkbh5* depletion induces G2/M cell cycle arrest, and impairs the MET process. **(A)** Cell proliferation was assessed by FACS measured by BrdU incorporation on day 3 of reprogramming using either scrambled shRNA or two different shRNAs targeting *Alkbh5*. **(B)** Quantification of mean percentage of each of the populations G1, S and G2/M from FACS data shown in figure 2A. The mean percentage of each population was written mean± S.D. **(C)** Cell proliferation was assessed by counting *Alkbh5*^*f/f*^ cells with or without addition of 1µM of 4-OH Tam for *Alkbh5* depletion. **(D)** Expression of mitotic checkpoint complex (MCC) factors as assessed by qPCR on day 3 of reprogramming using either scrambled shRNA or two different shRNAs targeting Alkbh5. The data are normalized to the housekeeping gene *Gapdh*. **(E)** Immunoblot analysis of protein level for several cell cycle regulators on day 3 of reprogramming using either scrambled shRNA or two different shRNAs targeting *Alkbh5*, and (A-TUB) used as loading control. **(F)** Expression of mesenchymal and epithelial genes as assessed by qPCR, on day 6 of reprogramming after infection either with scrambled shRNA or two different shRNAs targeting *Alkbh5*. The data are normalized to the housekeeping gene *Gapdh*. **(G)** Immunoblot analysis of protein level for mesenchymal and epithelial markers on day 6 of reprogramming after infection either with scrambled shRNA or two different shRNAs targeting *Alkbh5*, and A-TUB was used as loading control. Data are shown as mean ± SD; n = 3, *P < 0.05, **P < 0.01, and ***P < 0.001.

### ALKBH5 overexpression in the late phase enhances reprogramming efficiency by upregulating *Nanog*

We assessed the impact of ALKBH5 overexpression on the reprogramming process. Our data revealed that, overexpression of either ALKBH5, or ALKBH5 with a carboxyl terminal hemagglutinin (HA) tag (ALKBH5-HA), enhance the reprogramming process as measured by an increase in the percentage of SSEA1 positive cells and increase in number of ALP positive colonies at day 14 of reprogramming (Fig. 3A-C).

**Figure 3.**
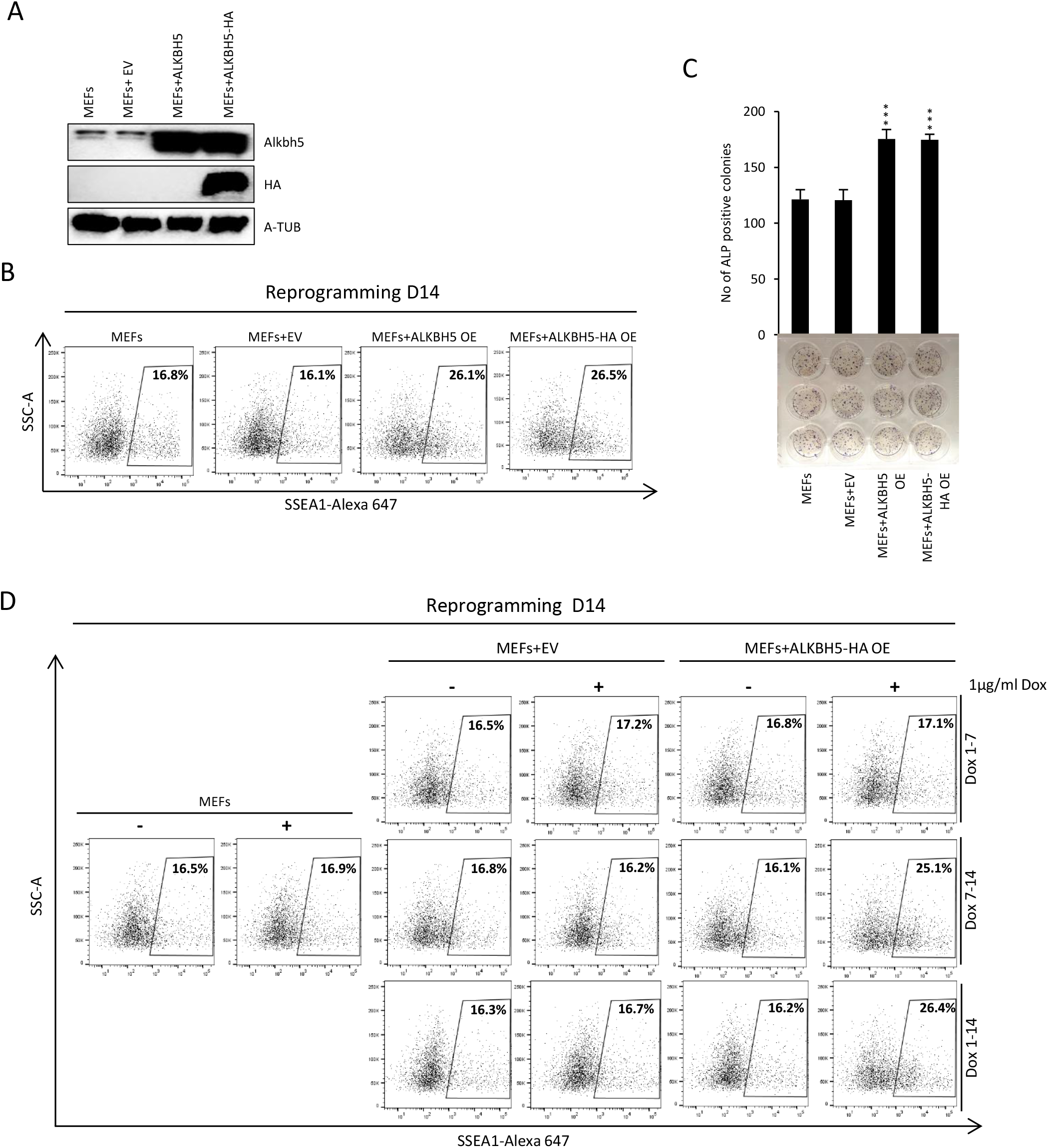
ALKBH5 overexpression enhances the reprogramming efficiency. **(A)** Immunoblot analysis of ALKBH5 protein level after lentiviral infection of MEFs with empty vector or ALKBH5 or ALKBH5 tagged with HA (ALKBH5-HA), and A-TUB used as loading control. **(B)** Fraction of SSEA1 positive cells determined by FACS analysis on day 14 of reprogramming after ALKBH5 overexpression. **(C)** Reprogramming efficiency was measured by counting the number of ALP positive colonies on day 14 of reprogramming after ALKBH5 overexpression. **(D)** Fraction of SSEA1 positive cells determined by FACS analysis on day 14 of reprogramming after temporal overexpression of ALKBH5-HA by 1µg/ml doxycycline (Dox) from day 1 to day 7, day 7 to day 14 or day 1 to day 14. MEFs were used as a negative control. Data are shown as mean ± SD; n = 3, *P < 0.05, **P < 0.01, and ***P < 0.001.

To investigate at what time of the reprogramming that ALKBH5 overexpression enhances reprogramming efficiency, we used a dox inducible overexpression system. We did not find any significant effect of ALKBH5 overexpression on the reprogramming efficiency at the early phase from day 1 to day 7. However, the percentage of SSEA1 positive cells at day 14 is greatly increased after overexpression of ALKBH5-HA from day 1-14, as well as after overexpression from day 7-14 only (Fig. 3D). To investigate the molecular mechanism responsible for enhancing reprogramming efficiency by overexpression of ALKBH5 at the late phase, we used a dox inducible system for temporal overexpression of ALKBH5 from day 10 to day 12 (Fig.4 A). We found that overexpression of ALKBH5 result in upregulation of the endogenous RNA level of reprogramming factors such as *Oct4, Sox2* and *Klf4*, and also other pluripotency factors including *Klf2, Tbx3, Rex1, Esrrb*, and in particular *Nanog* (Fig. 4B, C). We obtained similar results by overexpression of ALKBH5 from day 8 to day 10 (Supplementary Fig. 4 A-C). Previous studies have reported that *Nanog* is regulated postranscriptionally in both mouse and human ESCs by the m^6^A machinery [34, 35]. We hypothesized that *Nanog* transcripts are posttranscriptionally regulated through the m^6^A modification during reprogramming and that overexpression of the m^6^A demethylases ALKBH5 will reduce m^6^A levels, potentially affecting the stability of *Nanog* transcripts. To test this hypothesis in the reprogramming context, we did m^6^A IP at day 12 of reprogramming and indeed found that overexpression of ALKBH5 decreases the m^6^A level at *Nanog* transcripts (Fig.4D). Then by checking the stability of *Nanog* transcript after overexpression of ALKBH5, we found that overexpression of ALKBH5 increases the stability of *Nanog* transcripts (Fig.4E). Finally, we assessed whether ALKBH5 overexpression could rescue the *Alkbh5* KO phenotype in reprogramming. Our data revealed that overexpressing either ALKBH5 or ALKBH5-HA in *Alkbh5* KO MEFs could restore the reprogramming efficiency (Fig.4F supplementary Fig. D-F). Taken together, our findings suggest that ALKBH5 overexpression in the late phase of reprogramming enhances the reprogramming efficiency by decreasing the m^6^A level at *Nanog* transcripts, thus stabilizing these transcripts resulting in upregulation of *Nanog*.

**Figure 4.**
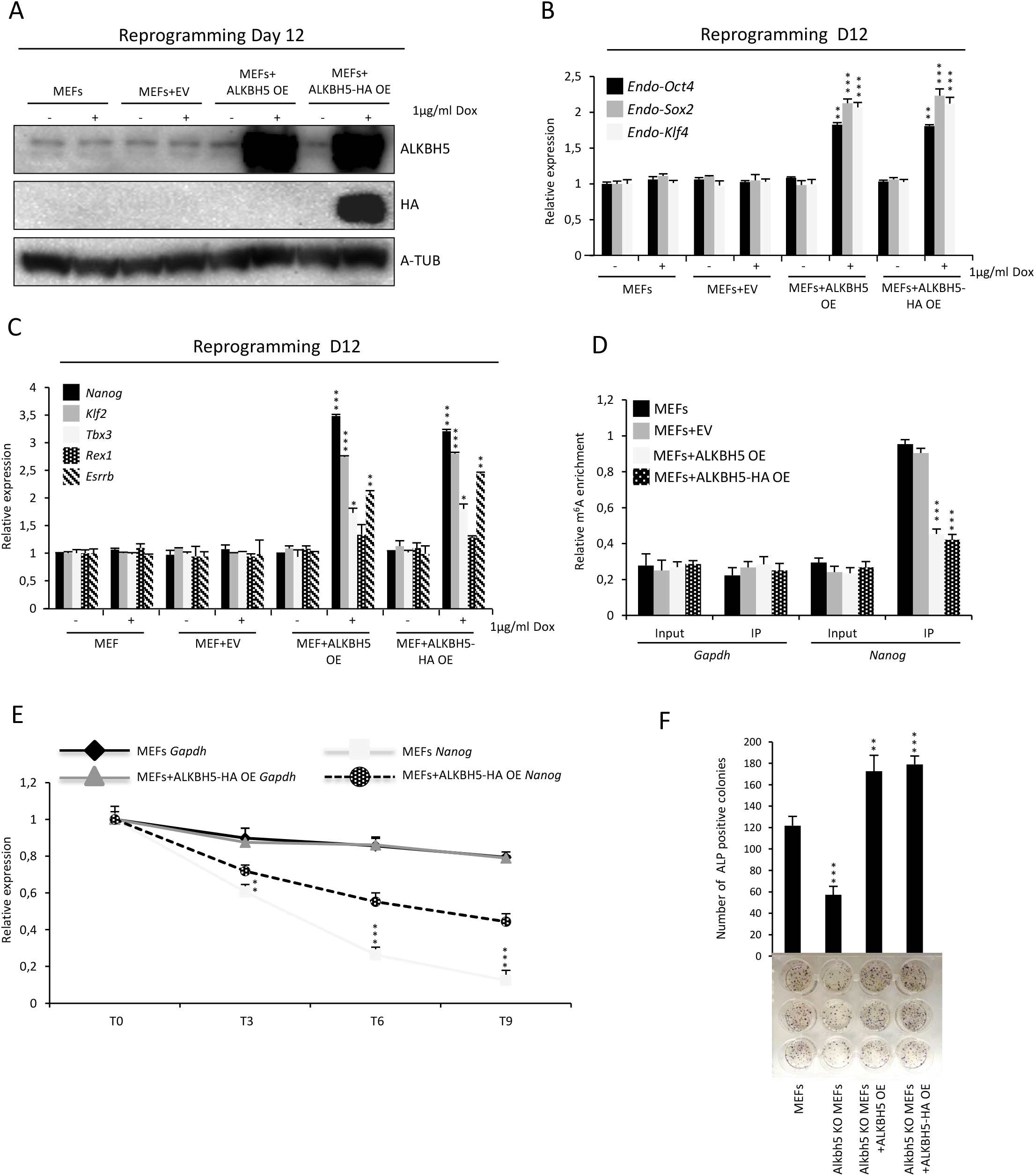
ALKBH5 overexpression in the late phase of reprogramming stabilizes *Nanog* transcripts resulting in increased *Nanog* expression. **(A)** Immunoblot analysis of ALKBH5 protein level after lentiviral infection of reprogrammed MEFs on day 12, with empty vector, ALKBH5 or ALKBH5-HA. 1µg/ml of Dox was added on day 10, then cells were harvested on day 12, and A-TUB used as loading control. **(B)** Endogenous expression of pluripotency factor (*Oct4, Sox2, Klf4*) as detected by qPCR, on day 12 of reprogramming using either empty vector, ALKBH5 or ALKBH5-HA. The data are normalized to the housekeeping gene *Gapdh*. **(C)** Expression of pluripotency markers detected by qPCR, on day 12 of reprogramming using either empty vector, ALKBH5 or ALKBH5-HA. The data are normalized to the housekeeping gene *Gapdh*. **(D)** m^6^A -IP qPCR data of Nanog on day 12 of reprogramming using empty vector, ALKBH5 or ALKBH5-HA overexpression. Data were shown as relative enrichment of m^6^A normalized to percentage (%) of input. **(E)** Half life time of *Nanog* mRNA on day of reprogramming after ALKBH5-HA. *Gapdh* was used a negative control, and data of cells treated with 5 µM Actinomycin D (ActD) were normalized to DMSO treated cells. **(F)** Reprogramming efficiency was measured by counting the number of ALP positive colonies on day 14 of reprogramming. Data are shown as mean ± SD; n = 3, *P < 0.05, **P < 0.01, and ***P < 0.001..

## Discussion

Ectopic expression of the four transcription factors OCT4, SOX2, KLF4, and c-MYC in somatic cells can establish the pluripotency regulatory circuitry, resulting in massive changes at both the epigenetic and transcriptional level and the generation of iPSCs [1, 2]. Successful therapeutic application of these iPSCs will likely require a comprehensive understanding of the molecular mechanism underlying somatic cell reprogramming. Here, we aimed to dissect the role of the m^6^A demethylases ALKBH5 in somatic cell reprogramming.

Resetting the pluripotency cell cycle pattern is an essential step of achieving successful iPSC generation, suggesting cell division rate is a key parameter for somatic cell reprogramming [36]. In agreement with that, *p53* and *Ink4/Arf* have been shown to act asbarriers to the reprogramming process [14-16]. Additionally, G2/M cell cycle regulators have been reported in maintaining pluripotency and the Cdk1/Cyclin B1 complex has been reported in enhancing the reprogramming process [37, 38]. Moreover, the m^6^A machinery has been reported to be involved in regulating Cdk1 and Cyclin B2, and knockout of Fat mass and obesity-associated (*Fto*) results in decreased expression of Cdk1 and Cyclin B2 causing G2/M cell cycle arrest in spermatogonia [39]. Here, we showed that *Alkbh5* depletion in MEFs or during early phase of somatic cell reprogramming decreased the expression of Cyclin B1 and B2 accompanied by cell cycle arrest at G2/M phase, which in turn resulted in reduced proliferation and MET transformation rate, ultimately leading to impaired reprogramming efficiency. We did not formally exclude the possibility that *Alkbh5* might have a direct effect on MET, which would be an interesting point for future studies. Furthermore, in contrast with that observed for the early phase of reprogramming, we found that depletion of *Alkbh5* in the late phase of reprogramming did not have a significant effect on reprogramming efficiency. This indicates that the negative effect *Alkbh5* depletion has on reprogramming efficiency plays out specifically during the early phase where both the resetting of the cell cycle pattern and morphological transformation to epithelial like cells occur.

Recent studies have revealed that the m^6^A modification on mRNA is essential in regulating pluripotency and self-renewal of stem cell, somatic cell reprogramming and early embryonic development [28-31]. Regulation of pluripotency by m^6^A has been reported in both mouse and human ESCs where *Mettl3* and/or *Mettl14* depletion induce a hyper-pluripotent state presumably through increasing the m^6^A level over several pluripotency related transcripts such as *Nanog*, resulting in increased transcript stability that hinder cells to exit from the pluripotency state [34, 35]. NANOG is a key regulator of pluripotency and is required for acquiring pluripotency during the late phase of reprogramming [40, 41]. A synergistic role of *Nanog* in overexpression together with DNA demethylating agents in the late phase of reprogramming has been reported to enhance acquisition of the pluripotency state [41, 42]. Moreover, NANOG co-binds with OCT4, SOX2 and KLF4 to many regulatory regions to facilitate binding of co-activator P300 [43]. Here we showed that ALKBH5 overexpression in the late phase of reprogramming decreases the m^6^A level at *Nanog* transcripts, resulting in increased *Nanog* stability leading to enhanced reprogramming efficiency. Consistent with our findings, ALKBH5 has been reported to positively regulate *Nanog* stability and expression in response to hypoxia-inducible factor (HIF)-1α- and HIF-2α in breast cancer stem cells (BCSCs) [31].

A recent study reported that YTHDF2/3, but not YTHDF1, regulate the MET event in somatic cell reprogramming in an m^6^A dependent manner through the Hippo signaling pathway effector *Tead2* [44]. Other studies have shown redundancy among the three paralogs *Ythdf1/2/3*, suggesting they can have adequate functional compensation at least in some biological contexts [45, 46]. It would be interesting to assess the role of *Ytdhf1/2/3*, as well as any redundancy, in the context of *Alkbh5* depletion in future studies.

In conclusion, we provide mechanistic insight into epitranscriptional regulation of somatic cell reprogramming by elucidating the biphasic regulatory role of ALKBH5 in modulating reprogramming efficiency at the posttranscriptional level in a stage specific manner (Fig. 5).

**Figure 5.**
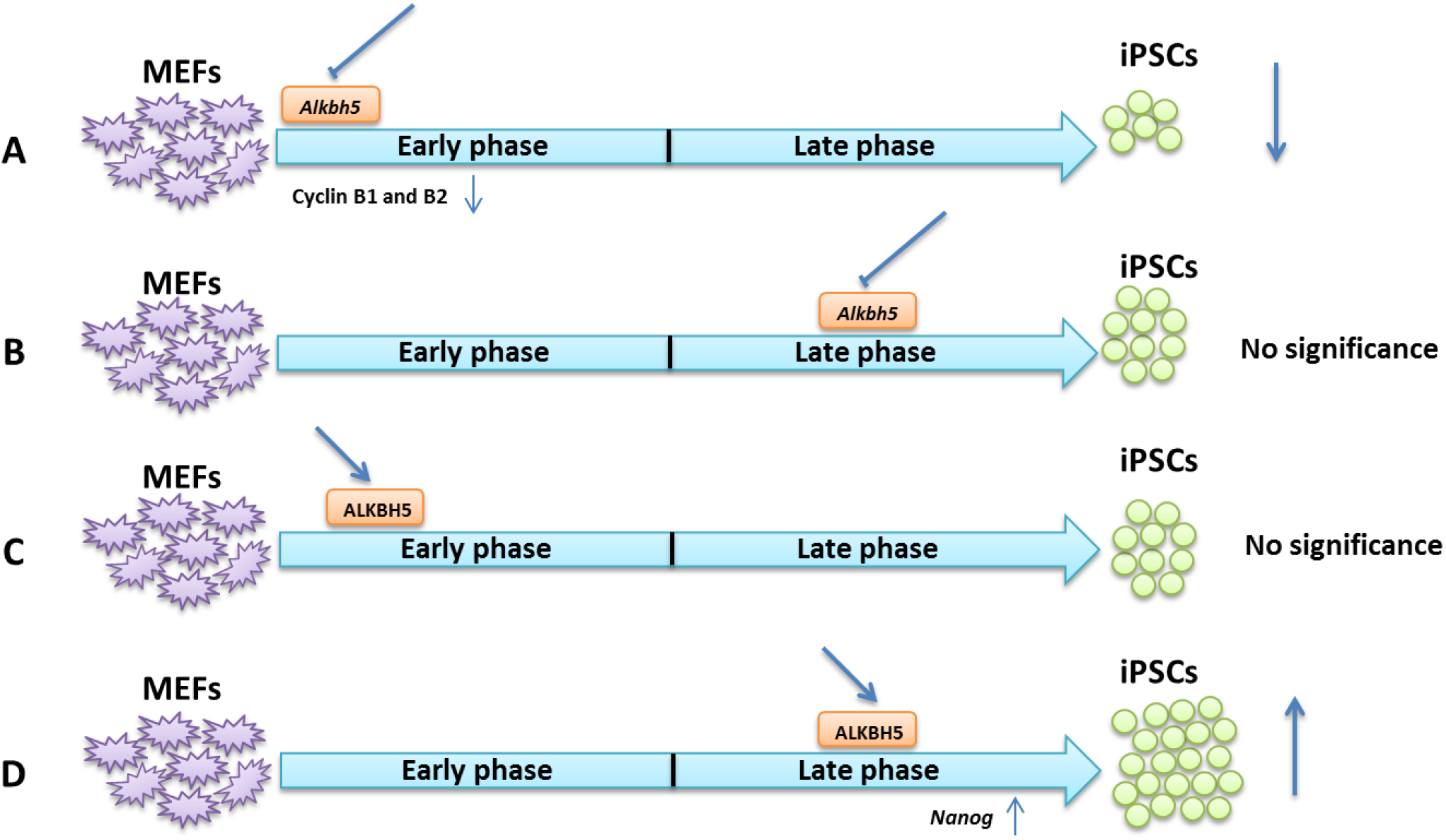
Model showing the biphasic role of ALKBH5 in somatic cell reprogramming. **(A**) Depletion of *Alkbh5* specifically in the early phase of reprogramming decreases the reprogramming efficiency by reducing the expression of cyclin B1 and B2. (**B)** Depletion of *Alkbh5* in the late phase of reprogramming has no impact on reprogramming efficiency. **(C)** Overexpression of ALKBH5 in the early phase of reprogramming does not affect the reprogramming efficiency. **(D)** Overexpression of ALKBH5 in the late phase enhances the reprogramming efficiency through increasing *Nanog* expression.

## Materials and Methods

### MEFs derivation

All of Wild type (WT), Knockout (KO) *Alkbh5* and homozygous floxed *Alkbh5* MEFs (*Alkbh5*^*f*/f)^ were derived from embryos at 13.5 d.p.c. Mice were housed and in Norwegian Transgenic Center (NTS). Briefly, pregnant mouse females were sacrificed on 13.5 or and embryos were dissected. The internal organs, head, and limbs were removed and used for genotyping. Then the remaining tissues were trypsinized using 0.25% trypsin for 30 min at 37°C with shaking to make single cell suspensions, then cells were pooled and plated in MEFs media until 80% confluence then trypsinized and stored in freezing solution (FBS+10%DMSO) in liquid nitrogen for future use. MEFs were cultured and maintained in DMEM+10% (tetracycline free FBS PAN-Biotech Catalog # P30-2602TC) till reaching to 70% to 80% confluence, then passaged at 1×10^5^ cells per well of 6-well plate.

*Alkbh5*^*f*/f^ MEFs were plated at 1×10^5^ cells per well of 6-well plate O.N, next day the cells were transfected with KA1153_pPBCAG-MerCreMer-IN (Addgene Plasmid #124183) together with PBase, and PB-GAC-Puro (a kind gift from professor Hitoshi Niwa, Kumamoto University IMEG) using Lipofectamin 2000 (Invitrogen# 11668019) or Fugene 6 (Promega #E2691) for 5 hours, then the medium was changed. Next day, the cells were cultured with medium containing 2µg/ml of Puromycin (Fisher Scientific # A1113803) for 2 days, and then the cells were treated with 1µM of 4 Hydroxy Tamoxifen (4-OH Tam) (Merk#H7904-5MG) for depletion of *Alkbh5* at indicated time points.

### Reprogramming

For reprogramming MEFs at early passages were plated as single cells at 1×10^5^ per well of 6 well plate or 5-6 x10^5^/ 10 cm dish depending on the purpose of experiment. The cells were infected with equal ratio of the retroviruses expressing the four reprogramming factors (Oct4, Sox2, Klf4, and c-Myc), and incubated at 37°C for 8-12 hours with 8 µg/ml of polybrene, then the medium changes next day. For either knockdown or overexpression experiments during reprogramming, the MEFs were plated at 1×10^5^ per well of 6 well, and infected with lentivirus for 8 hours, them medium was changed, and next day the selectable markers were added for 2 days. If the cells were trypsinized at day 7 reprogramming, the reprogrammed cells cultured with feeder layer CF-1 MEFs Irradiated, P3 2M (AMS biotechnology #GSC-6201G 2M or #GSC-6101G 7M) and LIF ESGRO® Recombinant Mouse LIF Protein (1000 units/mL) (Millipore # ESG1107). For induction of the transgene Stemolecule Doxycycline hyclate 10 mg (Stemgent#04-0016) was added at 1µg/ml every 2 days.

### Retrovirus preparation

Plate E cells were used for preparation of retrovirus (Cell bio labs #RV-101) Plate E cells were plated at 1×106 cells per 10 cm dish in DMEM%10FBS (tetracycline free FBS PAN-Biotech Catalog # P30-2602TC) till reaching to 70% to 80% confluence, then cells were transfected with 9 µg of each of pMXs-Oct4 (Addgene Plasmid #13366), pMXs-Sox2 (Addgene Plasmid #13367), pMXs-Klf4 (Addgene Plasmid #13370), pMXs-c-Myc (Addgene Plasmid #13375) per 10 cm dish using Fugene 6 (Catalog# Promega# E2691), and the medium was changed after 8 hours using (IMEDM+10%FBS). Retroviral supernatant were harvested after 48 and 72 hours, spin down at 1200 r.p.m for 5 minutes at 4 c, and then used freshly or frozen in aliquots at -80c. The viral titer was estimated to produce up to 7-8% SSEA1 at day 7 of reprogramming or using GFP control estimated more than 85% infection efficiency by FACS.

### Lentivirus preparation

LentiX 293T cells were used for preparation of lentivirus (Takahara Clontech #632180). Lentix 293T cells were plated at 1×10^6^ cells per 10 cm dish in DMEM%10FBS (tetracycline free FBS PANSera catalog) till the cells reaching to 70% to 80% confluence. Then cells were transfected with PsPAx2 (Addgene Plasmid #12260), and pMD2.G (Addgene Plasmid #12259), and the vector encoding either shRNA for Knockdown or overexpression Alkbh5 or Alkbh5-HA tagged at c terminal for overexpression using Fugene 6, and the medium was changed after 8 hours using (IMEDM+10%FBS). Retroviral supernatant were harvested after 48 and 72 hours, spin down at 1200 r.p.m for 5 minutes at 4 c, and then used freshly or frozen in concentrated form using (aliquots at -80c. The viral titer was estimated to produce up to 7-8% SSEA1 at day 7 of reprogramming.

### Cell proliferation assay

Mouse embryonic fibroblasts (MEFs) were plated at 1×10^4^ per well of 24 well plate at quadruplicate. Then, at each indicated time point four wells were trypsinized and counted independently using (Life Technologies #C10228 Countess™ Cell Counting Chamber Slides). Medium was replaced every 2 days and the data are presented as mean±SD for quadruplicate samples.

For reprogramming experiment, MEFs were plated at 1×105 cells per well of 6-well plate in triplicate, and infected with equal molar ratio of retroviral titer encoding Oct4, Sox2, Klf4, and c-Myc for 6 hours then medium changed, 8 hours after infection, cells were treated with either ethanol or 1µM of 4-hydroxy-Tamoxifen (4-OH-Tam) for depletion of Alkbh5. Cells were trypsinized at indicated time points and counted. Medium was replaced every 2 days and the data are presented as mean±SD for triplicate samples.

### Genotyping

Cells of tissue biopsies has been suspended in lysis buffer (1M Tris-PH 8, 5M NaCl, 0.5M EDTA PH8, 10% SDS) and freshly added (Proteinase K 20mg/ml) and incubated at 37c for 4hrs to O.N, then 300µl of 5M NaCl, then vortex, and incubated in ice for 10 min, then spinning at low speed, remove the supernatant, and transfer to new tube, then add 650µl Iso-propanol, vortex, and incubate at RT for 15 min, then centrifuge at 150,000 r.p.m, then discard the supernatant, and dissolve the pellet in 200µl TE buffer, then incubate at 55c for 10 min, then the DNA concentration is measured and 10-50 ng used per reaction.

### Cloning

Both mAlkbh5 and mAlkbh5-HAtag were amplified from the cDNA using gateway forward and reverse primer using PrimeSTAR GXL DNA Polymerase (Takahara Clontech # R050A-TAK), and the PCR product was purified using QIAquick PCR Purification Kit (Qiagen #28106), then shuttled to Gateway™ pDONR™221 Vector (Invitrogen#12536017) using Gateway™ BP Clonase™ II Enzyme mix (Invitrogen#11789020), then transformed to One Shot™ Stbl3™ Chemically Competent E. coli (Thermo Fisher #C737303), and positive clones were screen by colony PCR, and restriction digestion, and positive colonies were sent for sequencing. The correct clone was used as entry clone and then the cloned gene was shuttled to destination vector pLX301 (Addgene Plasmid #25895) For constitutive overexpression of either Alkbh5 or Alkbh5-HA, and pCW57.1 (Addgene Plasmid #41393) for periodic overexpression of either Alkbh5 or Alkbh5-HA using LR clonase (Thermo #11791020) based on manufacture protocol, and transformed to Stbl3 competent cells in case of Lentivirus destination vector, and positive clones were screen by colony PCR, and restriction digestion, and positive colonies were sent for sequencing, and confirmation, the positive colony was propagated and the plasmids were purified using Qiagen (Endotoxin free kit #12362), and used for making the virus.

#### For shRNA cloning

Two short hairpins shRNA for targeting mAlkbh5 were annealed in annealing buffer by heating 10 minutes at 95C In PCR machine then cooling by gradual decreasing the temperature to 4C in 30 minutes, and then the annealed oligos were ligated using T4 DNA Ligase (5 U/µL) (Thermo Fisher Scientific #EL0011) to either pLKO.1 puro (Addgene Plasmid #8453) for constitutive knockdown or Tet-pLKO-puro (Addgene Plasmid #21915) for periodic knockdown which was pre-linearized with AgeI-HF (NEB # R3552L) and EcoRI-HF (NEB#R3101S) restriction enzymes, then transformed to One Shot™ Stbl3™ Chemically Competent E. coli (Thermo Fisher #C737303), and several colonies were picked up and sent for sequencing. The positive clones were propagated and the plasmids were purified using Qiagen (Endotoxin free kit #12362), and used for making the virus.

### qPCR

TRIzol™ LS Reagent (Thermo scientific 10296010) was used for RNA extraction according to the manufacturer protocol, then the RNA was dissolved in UltraPure™ DNase/RNase-Free Distilled Water (Thermo Scientific 10977049), then 1µg was used to make the cDNA using SuperScript™ IV VILO™ Master Mix with ezDNase™ Enzyme (Thermo Scientific 11766050) based on manufacturer protocol. For Real time PCR, 2µl of cDNA was used per reaction using PowerUp™ SYBR™ Green Master Mix (Thermo Scientific A25777). The transcript level was normalized to the internal control. List of primers is attached in supplementary table 1

### RNA stability

Cells were treated with 5µg/ml of actinomycin D (Tocris #1229). At indicted time point 3, 6, and 9 hours total RNA was extracted and DMSO treated cells was used as a control, and relative RNA expression was detected by qPCR

### m6A dot blot

Total RNA was extracted from cells using TRIzol™ LS Reagent (Thermo scientific 10296010) or RNeasy Plus Mini Kit (Qiagen# 74134). mRNA was isolated and purified using Dynabeads™ mRNA Purification Kit (for mRNA purification from total RNA preps) (Invitrogen # 61006) following the manufacturer’s instructions. For m6A dot blot, mRNA was hybridized onto the Hybond-N+ membrane (GE Healthcare). After crosslinking spotted mRNA to membrane using Stratalinker 2400 UV Crosslinker, the membrane was blocked with 5% skimmed milk for 1 h, incubated with mouse anti-m6A antibody (1:1000, Millipore # MABE1006) at 4°C overnight. Then the membrane was incubated with HRP-conjugated donkey anti-mouse IgG at room temperature for 1 h. The membrane was photographed using the ECL imaging system (Bio-Rad). Finally, the membrane was stained with 0.02% methylene blue. Relative m6A level was quantified using ImageJ.

### m6A IP-qPCR

Control and Alkbh5-HA overexpressed reprogrammed cells at day 12 of reprogrammed were harvested and mRNA was extracted from RNA as described previously. 1 to 2 µg of mRNA was fragmented for 4 minutes at 70°c for 4 minutes, and then mRNA was precipitated and the pellet was dissolved in Ultrapure DNase/RNase free water, then incubated with pre conjugated m6A/protein G (Dynabeads™ Protein G for Immunoprecipitation#10003) beads in IP buffer, and incubated at 4c for O.N. The mRNA was isolated from the beads using Trizol LS, and the RNA was used to make cDNA using SuperScript™ IV VILO™ Master Mix with ezDNase™ Enzyme (Thermo Scientific 11766050) based on manufacturer protocol. The m6A mRNA level was finally determined by real-time quantitative PCR relative to the input.

### Western blot

Cells were washed twice with ice cold 1xPBS, and then scrapped and transferred to 1.5 ml Eppendorf tube, then centrifuged, and supernatant was discarded, and the cells were lysed on RIPA lysis buffer (20mM Tris-HCl PH7.5, 1mM MgCl2, 500mM NaCl, 20% glycerol, 0.5% NP-40, 1mM EDTA, 1mM EGTA) and freshly added 1x Halt Protease Inhibitor cocktail (100X) (Thermo Fisher #87786), then incubated on ice for 30 min, then centrifuged at maximum speed for 30 minutes, then the supernatant was transferred to new Eppendorf, and then the protein content was measured using Bradford protein assay (BSA) method, and then equal amounts of protein was lysed with 1x Bolt™ LDS Sample Buffer (Thermo Scientific B0008), and 1x Bolt™ Sample Reducing Agent(Thermo Scientific B0009), and loaded on Bolt ready gel (4-12%), and then the protein was transferred to PVDF or nitrocellulose Biorad pads using Trans-Blot Turbo Transfer System. Then the membrane was blocked using 5% skimmed milk in 1xTBST buffer, and then incubated with the primary antibody O.N was agitation. Next day, the membrane was washed 3 times using 1xTBST buffer, then incubated with the secondary antibody for 1 hr at RT, and then washed 3 times using 1xTBST buffer, and then the protein detected with Pierce™ ECL Western Blotting Substrate (Thermo Fisher 32209) or SuperSignal™ West Femto Maximum Sensitivity Substrate (Thermo Fisher 34094), using Biorad ChemiDoc XRS, and Precision Plus Protein™ Dual Color Standards (Bio-Rad#161-0374) as a protein standard. Antibodies list is attached supplementary table 2.

### Alkaline phosphatase staining

Alkaline phosphatase staining has been already done using Leukocyte Alkaline Phosphatase Kit (Sigma 85L3R), based on the manufacturer protocol as previously described (Khodeer and Era, 2017).

### Cell cycle analysis

The cells were trypsinized, harvested and collected, and washed 2 times with 1xPBS, and then suspended in 300µl ice cold 1xPBS, and then 700µl ice cold 100% ethanol and incubated at 4°C for at least 30 minutes. Then the cells centrifuged and supernatant was aspired and suspended in 200µl (Propidium Iodide (PI)/RNase Staining Solution, (Cell signaling 4087S) and incubated at RT for 30 minutes and then analyzed by FACS.

### Apoptosis

Detection of apoptotic cells was done by using FITC-Annexin V Apoptosis Detection Kit with 7-AAD (Biolegend#640922). Briefly, The cells were collected, and washed 2 times with 1xPBS, and then suspended in 200µ 1x binding buffer, 1µl Annexin V-FITC, and 7AAD (1:200) and incubated at RT for 30 minutes in the dark. The cells were centrifuged and suspended in 300µl 1x binding buffer, and then analyzed by FACS.

### SSEA1 staining

Cells at indicated time points were washed two times with 1xPBS, and trypsinized, precipitated, and counted. Then 1×106 cells were washed again with 1x Hanks buffer and stained with 5µl of Alexa Fluor® 647 anti-mouse/human CD15 (SSEA-1) Antibody (Biolegend#125608) in100µl BD Pharmingen™ Stain Buffer (FBS) (BD Biosciences #554656) for 30 min on Ice, and then cells were washed once with 1x Hanks buffer, then 7AAD (1:200), and SSEA1 positive fraction was analyzed using FACS BD Fortessa.

### BrdU incorporation assay

APC BrdU Flow Kit (BD Biosciences #552598) was used according to the manufacturer’s protocol. Briefly, cells were labeled by adding 10 μM of BrdU to the culture medium. Treatment was done for 1 hour and then cells were fixed and permeabilized, and then treated with DNase for 1 hour at 37°c, then stained with anti-BrdU APC for 20 minutes at RT, then resuspended in 7AAD and analyzed by FACS.

### Statistical Analysis

All data were collected from at least three independent experiments. Data were analyzed using Student’s *t*-test or one-way ANOVA followed using Graphpad software. Significance was presented as ^*^*p* < 0.05, ^**^*p* < 0.01, and ^***^*p* < 0.001. Error bars represented mean±SD.

## Figure legends

**Supplementary figure 1.**
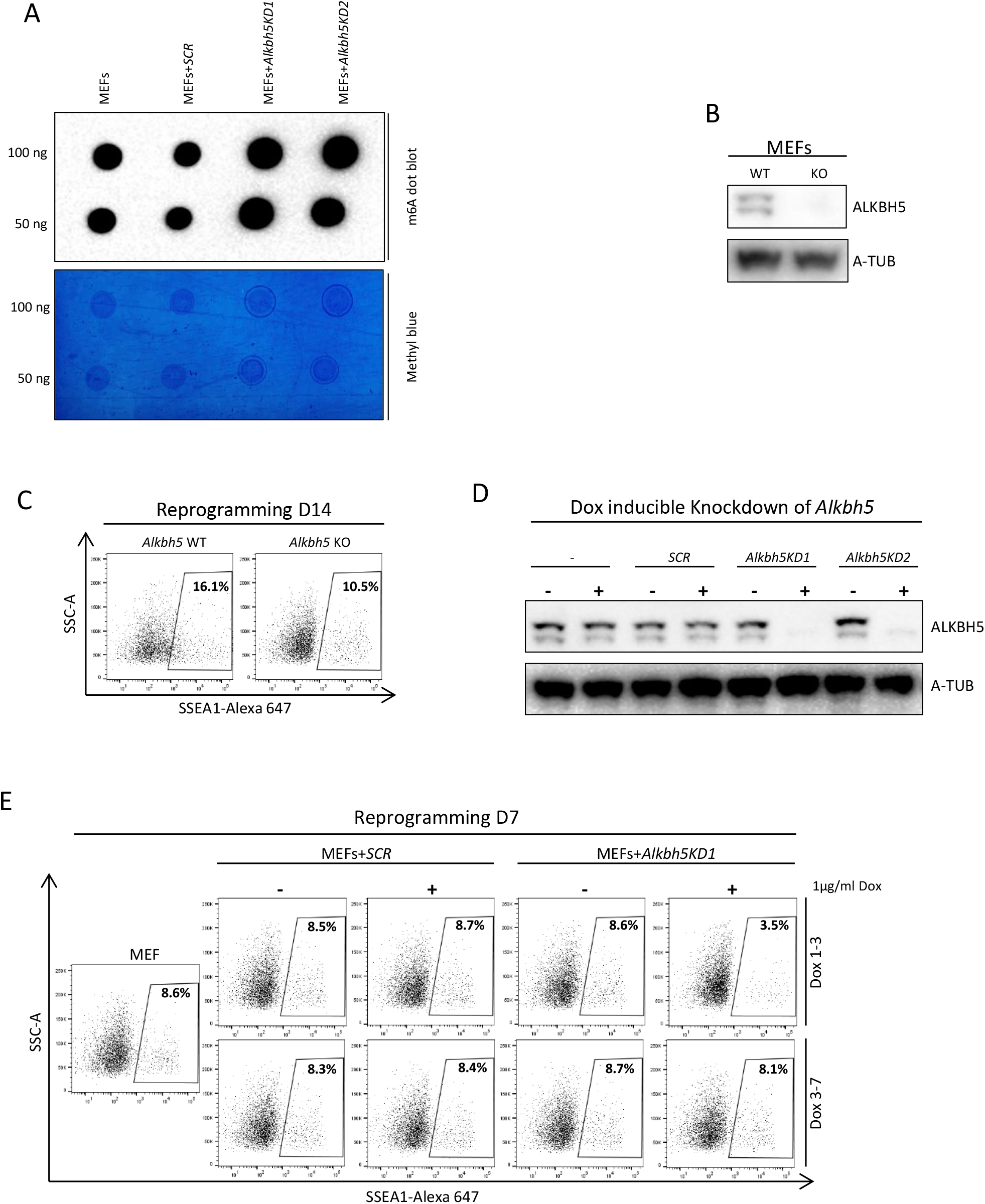
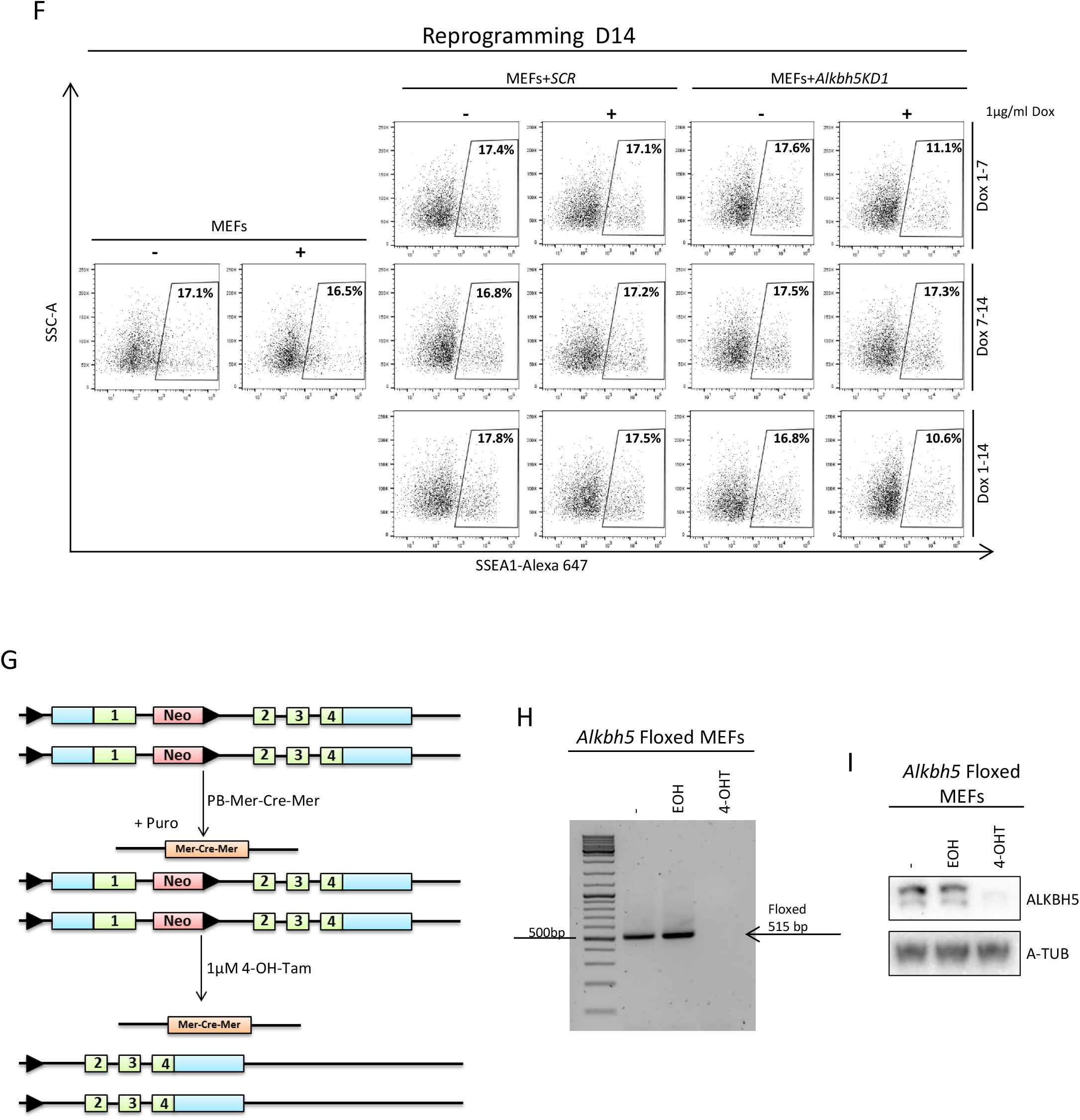
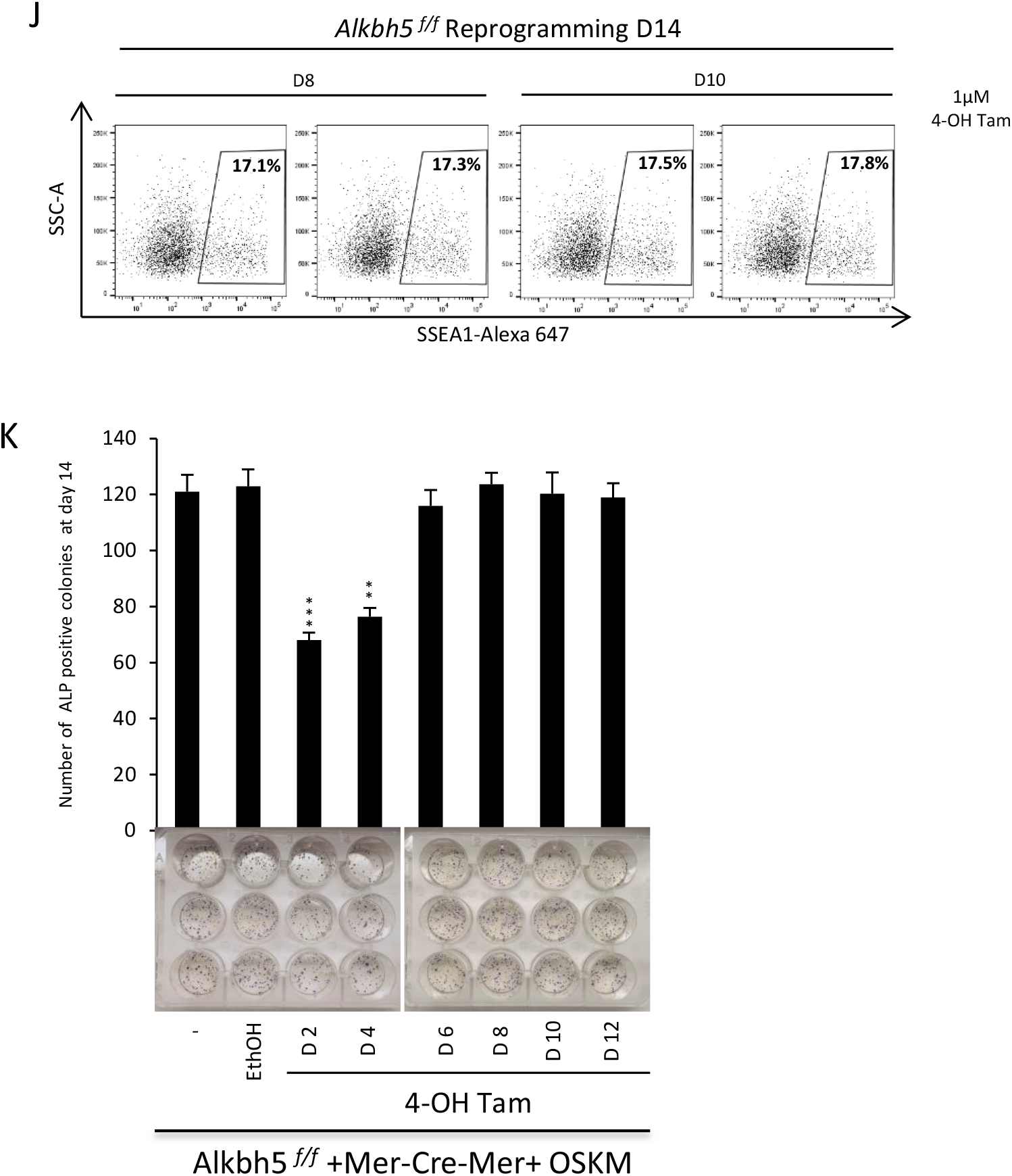
Depletion of *Alkbh5* in the early stage impairs the reprogramming efficiency. **(A)** m^6^A dot blot analysis of uninfected MEF or infected either with lentiviral encoding for scrambled shRNA, and two different shRNAs targeting *Alkbh5* (upper panel). Methyl blue staining was used as control to eliminate the difference in loaded mRNA amount (lower panel). **(B)** Immunoblot analysis of ALKBH5 protein level in WT and *Alkbh5* KO MEFs. A-TUB was used as loading control. **(C)** Fraction of SSEA1 positive cells determined by FACS in WT and *Alkbh5* KO reprogrammed MEFs on day 14 of reprogramming. **(D)** Immunoblot analysis of ALKBH5 protein level in MEFs infected with lentiviral encoding for scrambled shRNA, and two different shRNAs targeting *Alkbh5*. After selection with puromycin for 2 days, cells were treated with 1µg/ml of Dox to induce the expression of shRNA. A-TUB was used as loading control. **(E)** Fraction of SSEA1 positive cells determined by FACS in reprogrammed MEFs infected either by scrambled shRNA or shRNA targeting *Alkbh5* with or without 1µg/ml Dox treatment on day 7 of reprogramming. MEFs were used as a negative control. **(F)** Fraction of SSEA1 positive cells determined by FACS in reprogrammed MEFs infected either by scrambled shRNA or shRNA targeting *Alkbh5* with or without 1µg/ml Dox treatment on day 14 of reprogramming. MEFs were used as a negative control. **(G)** Experimental design for *Alkbh5* depletion. Homozygous *Alkbh5*^*f/f*^ MEFs were derived from mice at 13.5 days post coitum (d.p.c), before transfected with PB-GAG-Mer-Cre-Mer, then selection with puromycin for 2 days, and treatment with 1µM 4-OH Tam for induction of the Cre to remove *Alkbh5*. **(H)** Genotyping of homozygous *Alkbh5*^*f/f*^ MEFs untreated or treated with either ethanol (negative control) or 1µM 4-OH Tam for *Alkbh5* removal. The band corresponds to the Neomycin (Neo) PCR amplicon of 515 base pairs (bps). **(I)** ALKBH5 Immunoblot analysis of homozygous *Alkbh5*^*f/f*^ MEFs untreated or treated with either ethanol (negative control) or 1µM 4-OH Tam for *Alkbh5* removal. **(J)** Fraction of SSEA1 positive cells determined by FACS in reprogrammed MEF on day 14 of reprogramming. Reprogrammed homozygous *Alkbh5*^*f/f*^ MEFs treated with 1µM 4-OH Tam for *Alkbh5* depletion at day 8 or day 10 of reprogramming. **(K)** Reprogramming efficiency as assessed by counting the number of ALP positive colonies on day 14 of reprogramming. Reprogrammed homozygous *Alkbh5*^*f/f*^ MEFs treated with 1µM 4-OH Tam for *Alkbh5* at day 2, 4, 6, 8, 10 and 12 of reprogramming, and ethanol treatment was used as negative control. Data are shown as mean ± SD; n = 3, *P < 0.05, **P < 0.01, and ***P < 0.001.

**Supplementary figure 2.**
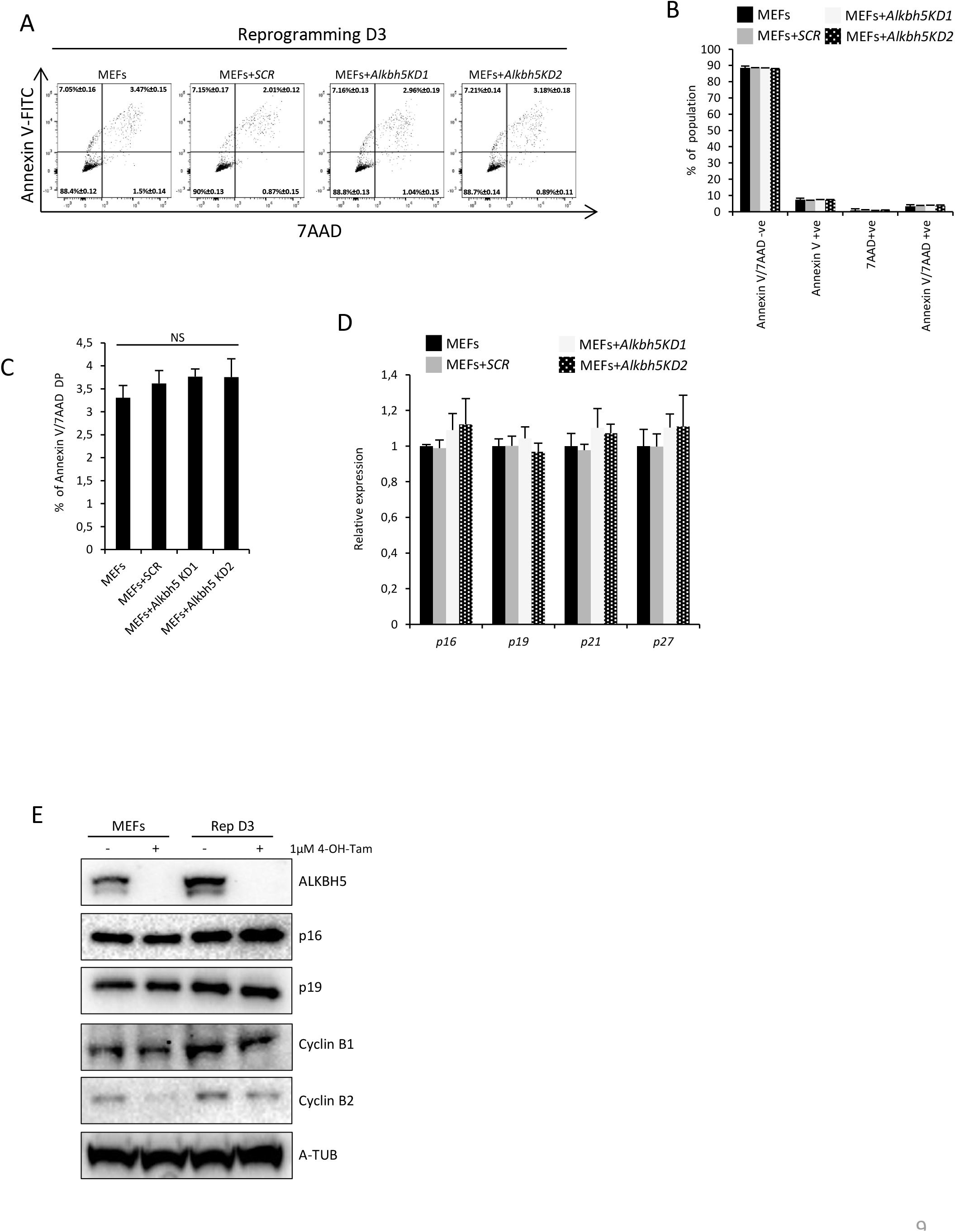
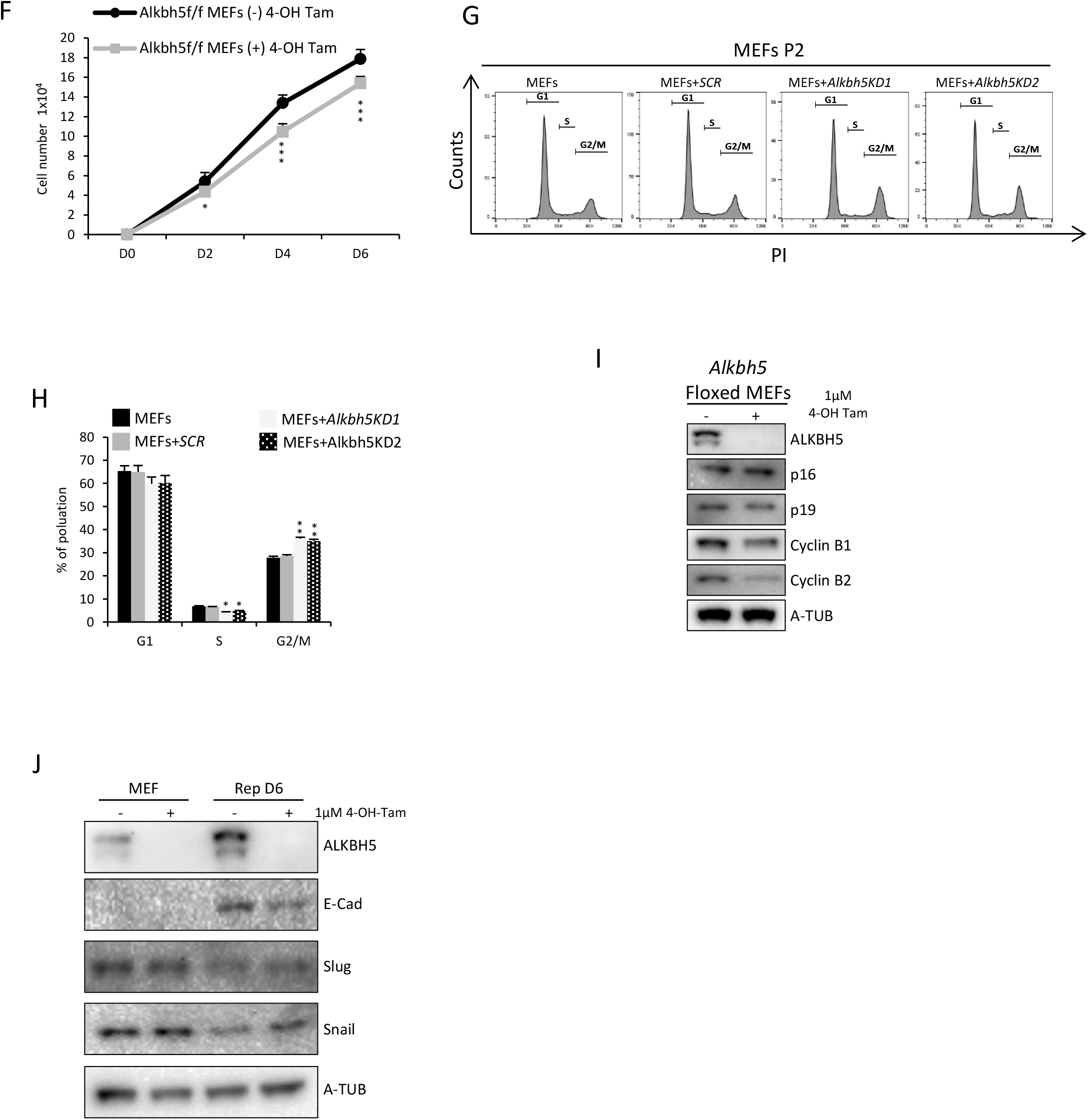
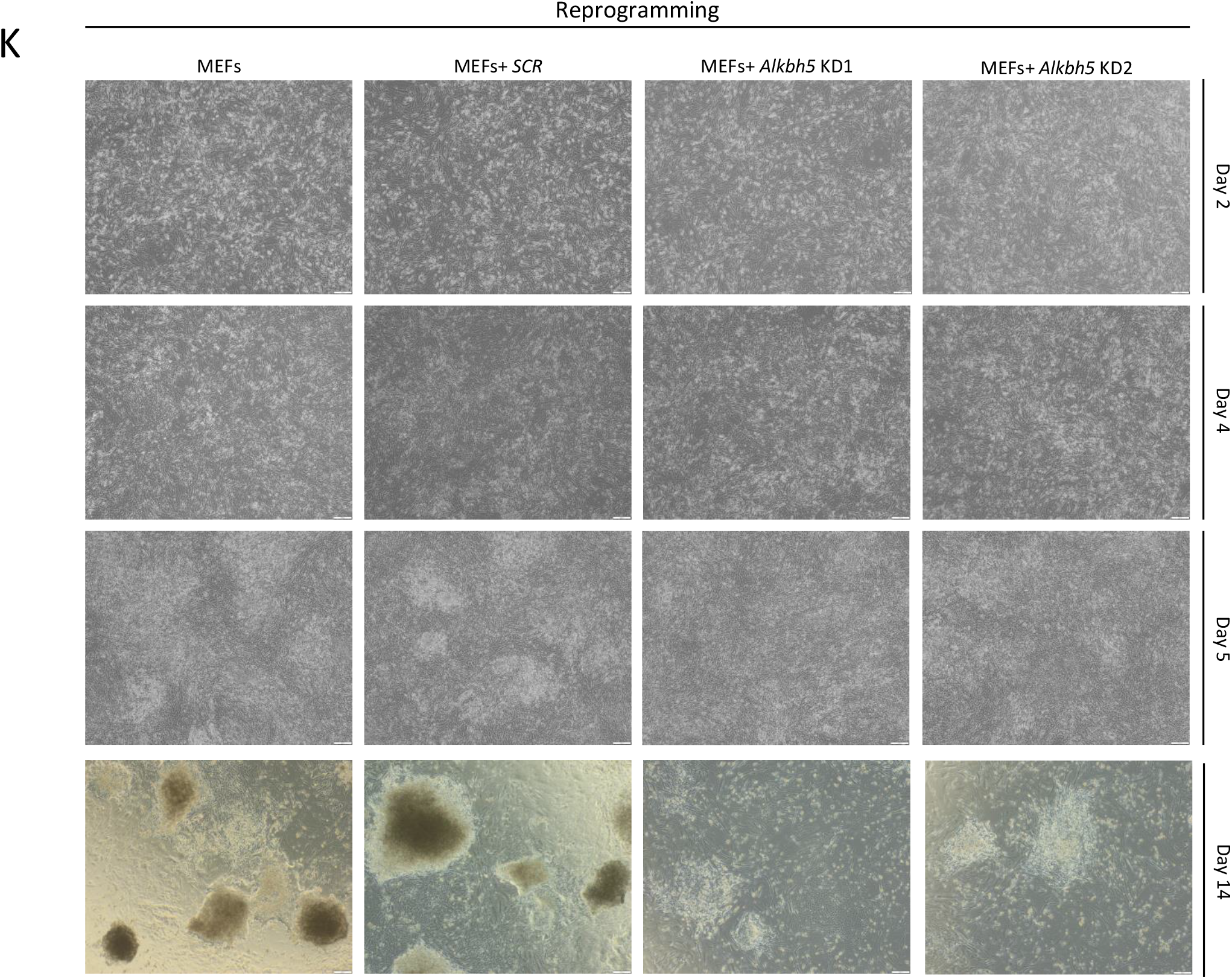
*Alkbh5* removal impairs the cell proliferation in either MEFs or reprogrammed MEF without increasing apoptosis. **(A)** Fraction of apoptotic cells determined by FACS in reprogrammed MEFs uninfected or infected either by scrambled shRNA or two shRNAs targeting *Alkbh5* was assessed at day 3 of reprogramming using double staining with Annexin V and 7AAD staining. **(B)** Analysis of cell apoptosis data determined by FACS in (Supplementary Fig. 2 A), each of 7AAD or Annexin V single positive (+ve) or negative (-ve), Annexin V/7AAD +ve or Annexin V/7AAD –ve. **(C)** Only Annexin V/7AAD double positive population from (Supplementary Fig. 2 B) to clarify insignificance among reprogrammed MEFs; uninfected or infected either by scrambled shRNA or two shRNAs targeting *Alkbh5* .N.S; Not significant. **(D)** Expression of G1 cell cycle regulators as assessed by qPCR at day 3 of reprogramming in reprogrammed MEFs uninfected or infected either by scrambled shRNA or two shRNAs targeting *Alkbh5* was estimated at day 3 of reprogramming. The data are normalized to the housekeeping gene *Gapdh*. **(E)** Immunoblot analysis of protein level for several cell cycle regulators in either homozygous *Alkbh5*^*f/f*^ MEFs or reprogrammed homozygous *Alkbh5*^*f/f*^ MEFs on day 3 with or without treatment with 1µM 4-OH Tam treatment to remove *Alkbh5*. A-TUB was used as loading control. **(F)** Cell proliferation assay homozygous *Alkbh5*^*f/f*^ MEFs with or without treatment with 1µM 4-OH Tam treatment to remove *Alkbh5* at different time points. **(G)** Cell cycle analysis detected by PI staining and analyzed by FACS in uninfected MEFs or infected with scrambled shRNA or two different shRNAs targeting *Alkbh5*. **(H)** Quantification of cell cycle phase G1, G2 and G2/M of data from Supplementary Fig. 2 G. **(I)** Immunoblot analysis of protein level for several cell cycle regulators in homozygous *Alkbh5*^*f/f*^ MEFs with or without treatment with 1µM 4-OH Tam treatment to remove *Alkbh5*. A-TUB used as loading control. **(J)** Immunoblot analysis of protein level for both mesenchymal and epithelial markers in either homozygous *Alkbh5*^*f/f*^ MEFs or reprogrammed homozygous *Alkbh5*^*f/f*^ MEFs on day 6 with or without treatment with 1µM 4-OH Tam treatment to remove *Alkbh5*. A-TUB used as loading control. **(K)** Phase contrast images of tracking morphological changes during reprogramming. Reprogrammed MEFs uninfected or infected either by scrambled shRNA or two shRNAs targeting *Alkbh5* was estimated at day 2, 4, 6 and 14 of reprogramming. Data are shown as mean ± SD; n = 3, *P < 0.05, **P < 0.01, and ***P < 0.001.

**Supplementary figure 4.**
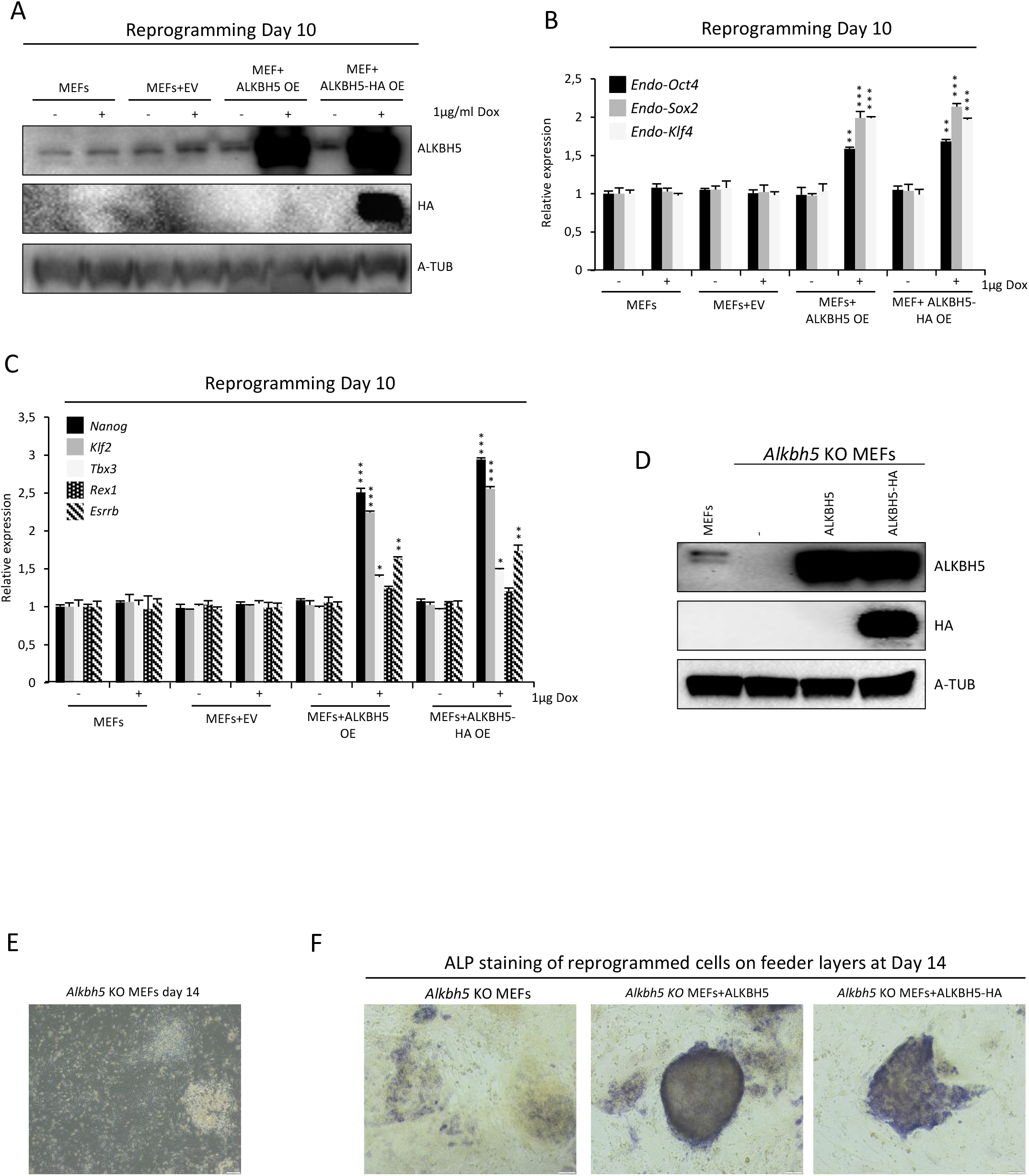
ALKBH5 overexpression in the late phase of reprogramming enhances the reprogramming efficiency through increasing *Nanog* expression. **(A)** Immunoblot analysis of ALKBH5 protein level after lentiviral infection of reprogrammed MEFs on day 12 with empty vector, ALKBH5 or ALKBH5-HA. 1µg/ml of Dox was added on day 8 then cells were harvested on day10. A-TUB used as loading control. **(B)** Endogenous expression of pluripotency factors (*Oct4, Sox2, Klf4*) as detected by qPCR, on day 10 of reprogramming using either empty vector, ALKBH5 or ALKBH5-HA. The data are normalized to the housekeeping gene *Gapdh*. **(C)** Expression of pluripotency markers detected by qPCR, on day 12 of reprogramming using either empty vector, ALKBH5 or ALKBH5-HA. The data are normalized to the housekeeping gene *Gapdh*. **(D)** Immunoblot of ALKBH5 in WT and KO *Alkbh5* MEFs, and rescued KO MEFs infected with lentiviral ALKBH5 and ALKBH5-HA. A-TUB used as loading control. **(E)** Phase contract image of *Alkbh5* KO reprogrammed MEFs at day 14 of reprogramming. **(F)** Phase contrast image of ALP stained of reprogrammed *Alkbh5* KO MEFs, and rescued KO MEFs infected with either ALKBH5 or ALKBH5-HA at day 14. Data are shown as mean ± SD; n = 3, *P < 0.05, **P < 0.01, and ***P < 0.001.

## Acknowledgments

The authors thank the Norwegian Transgenic Center (Norsk Transgen Senter (NTS)) and in particular; Shiasta Khan, Ingunn Jermstad, and Dr. Knut Tomas Dalen for setting up and maintaining floxed *Alkbh5* mice mating, Guro Flor Lien and Gaute Nesse for maintaining and genotyping knockout *Alkbh5* mice. This work was supported by the South?Eastern Norway Regional Health Authority Grant 2018063, and by the Research Council of Norway, Young Research Talents Grant 289467 (to John Arne Dahl).

